# Structure of the human UBR5 E3 ubiquitin ligase

**DOI:** 10.1101/2022.10.31.514604

**Authors:** Feng Wang, Qing He, Wenhu Zhan, Ziqi Yu, Efrat Finkin-Groner, Xiaojing Ma, Gang Lin, Huilin Li

**Author notes:** These authors contributed equally to this work (Feng Wang, Qing He).

## Abstract

The human UBR5 (also known as EDD) is a single polypeptide chain HECT-type E3 ubiquitin ligase essential for embryonic development in mammals. Although widely expressed, *UBR5* is markedly amplified and overexpressed in breast, ovarian, prostate, gastric and pancreatic cancers. Dysregulated UBR5 functions like an oncoprotein to promote cancer growth and metastasis, making UBR5 a potential target for therapeutics. Unexpectedly, we found that human UBR5 assembles a dimer and a tetramer in solution. We determined the dimer structure at 2.8 Å and the tetramer structure at 3.5 Å average resolution. UBR5 is a crescent shaped molecule with a seven-bladed β-propeller and two small β-barrel domains (SBB1/2) at the N-terminal region, a catalytic HECT domain at the C-terminus, and an extended helical scaffold and an N-degron-recognizing UBR box in the middle. The dimer is assembled as a stable head-to-tail dimer via extensive interactions in the middle helical scaffold region. The tetramer is assembled via SBB2-SBB2 interaction from two face-to-face dimers, forming a large cage with all four catalytic HECT domains facing the central cavity. Importantly, the N-terminal region of one subunit and the HECT of the other form an “intermolecular jaw” in the dimer. Using enzymatic and cellular assays, we showed that the jaw-lining residues are important for function, suggesting that the intermolar jaw functions to recruit ubiquitin loaded E2 to UBR5 for the transthiolation reaction. Further work is needed to understand how oligomerization regulates the UBR5 ligase activity. This work provides a framework for structure-based anticancer drug development against the distinctive HECT E3 ligase and contributes to a growing appreciation of E3 ligase diversity.

## INTRODUCTION

The Ubiquitin (Ub)–Proteasome System (UPS) is responsible for a large portion of regulated proteostasis in eukaryotes (Morreale and Walden, 2016). In this system, a cascade of three enzymes (the Ub-activating enzyme E1, Ub-conjugating E2, and Ub ligase E3) act sequentially to ubiquitylate target proteins, with the E3 predominantly determining the substrate specificity (Grabarczyk et al., 2021; Weissman, 2001). Most E3s can be classified into three families: The “Really Interesting New Gene” (RING) family, the “Ring-Between-Ring” (RBR) family, and the “Homology to E6AP C-Terminus” (HECT) family. The RING family members are often formed by multi-protein complexes and mediate direct transfer of Ub from an Ub-loaded E2 to a substrate without forming an E3–Ub intermediate (Duda et al., 2011; Petroski and Deshaies, 2005). In contrast, the RBR and HECT family members are usually single polypeptide chains and catalyze a two-step reaction in which Ub is first transferred from Ub–E2 to the E3 to form a Ub–E3 covalent intermediate, followed by a second transfer of the Ub to a substrate (Grabarczyk et al., 2021; Hunkeler et al., 2021; Lechtenberg et al., 2016; Rotin and Kumar, 2009; Walden and Rittinger, 2018). Humans encode 600-700 E3 ligases, accounting for ~5% of the genome (Berndsen and Wolberger, 2014; George et al., 2018; Li et al., 2008). Perhaps due to their giant size and often flexible architecture, only a dozen or so full-length E3 ligase structures have been reported (Horn-Ghetko and Schulman, 2022; Sherpa et al., 2022), including the yeast Ubr1 (Pan et al., 2021), the human and parasite HUWE1 (Grabarczyk et al., 2021; Hunkeler et al., 2021), the yeast GID3 complex (Qiao et al., 2020; Sherpa et al., 2021), the chicken Fanconi anemia core complex (Shakeel et al., 2019), the mouse AAA-E3 ligase RNF213 (Ahel et al., 2020), the human ASB9 (Lumpkin et al., 2020), the human BRCA1-BARD E3 ligase (Hu et al., 2021; Witus et al., 2021), and two human E3-E3 super-assemblies, SCF–ARIH1 and CUL5–ARIH2 (Horn-Ghetko et al., 2021; Kostrhon et al., 2021).

Humans encode seven N-end rule pathway E3 ligases, UBR1-7, four of which (UBR1, UBR2, UBR4 and UBR5) contain an N-degron recognition UBR Box (Kim et al., 2021). UBR5 is unique among them with a HECT domain and therefore is the only one that belongs to the HECT family (Sriram and Kwon, 2010). UBR5 is also known as EDD or EDD1 (E3 ligase identified by Differential Display) and HYD (hyperplastic discs) (Callaghan et al., 1998; Mansfield et al., 1994). UBR5 has a wide range of substrates, including β-catenin, TopBP1, TERT, CDK9, ATMIN, PEPCK1, CAPZA1, CDC73 (Xiang et al., 2022), which are implicated in multiple cellular processes, such as DNA damage, metabolism, transcription, apoptosis, and immunoregulation (Cojocaru et al., 2011; Hay-Koren et al., 2011; Honda et al., 2002; Rutz et al., 2015; Wang et al., 2013; Yoshida et al., 2006; Zhang et al., 2014), and UBR5 has been implicated in disassembly of mitotic checkpoint complexes (Kaisari et al., 2022). Hence, UBR5 is a key regulator of cell signaling relevant to broad areas of cancer biology (Shearer et al., 2015). In addition, UBR5 is highly expressed in a substantial proportion of breast, ovarian, prostate, gastric, and pancreatic cancers (Ding et al., 2020; Li et al., 2021; Liao et al., 2017; Song et al., 2020; Yang et al., 2020). Recently, UBR5 E3 pathway was shown to play a key role in the aggression of breast and ovarian cancers by enabling the cancer cells to survive and resist immune attack and standard treatments (Liao et al., 2017; Song et al., 2020; Xiang et al., 2022). The tumor-promoting effect of UBR5 has been largely attributed to its ubiquitin ligase activity, highlighting the potential for targeting UBR5 for anticancer drug development and the importance of determining UBR5’s structure and molecular mechanism (Song et al., 2020).

UBR5 contains 2799 amino acids and has an estimated mass of 309 kDa (**Fig. 1a**). The HECT domain is at the C-terminus with a larger N-lobe that interacts with E2s and a C-lobe containing the catalytic Cys2768 (Matta-Camacho et al., 2012). In addition to HECT, UBR5 also contains a Ub association domain (UBA) that interacts with Ub (Kozlov et al., 2007; Ohtake et al., 2018), two predicted small β-barrel domains (SBB1 and SBB2), a 70-residue zinc finger UBR-Box that recognizes the N-degron (Choi et al., 2010; Matta-Camacho et al., 2010; Sriram and Kwon, 2010), and an MLLE (the term comes from a signature motif, MLLEKITG) domain that mediates protein-protein interactions (Muñoz-Escobar et al., 2015). Structures of the C-lobe and MLLE have been reported (Matta-Camacho et al., 2012; Muñoz-Escobar et al., 2015). Additionally, previous studies have determined the crystal structures of the isolated HECT domains of several HECT E3 ligases (Kamadurai et al., 2013; Kamadurai et al., 2009; Lorenz, 2018; Maspero et al., 2011; Maspero et al., 2013; Ogunjimi et al., 2005; Singh et al., 2019). However, the structure of the full-length UBR5 has been lacking. We have determined the cryo-EM structure of the human full-length UBR5 at up to 2.66 Å resolution and revealed that UBR5 assembles stable dimers and tetramers in solution. The high-resolution structures shed light on the UBR5 catalyzed transthiolation reaction mechanism and provide a platform for developing UBR5 inhibitors for the potential treatment of cancers. Future studies are required to understand whether the UBR5 dimer and tetramer possess distinct functions or substrates.

**Figure 1.**
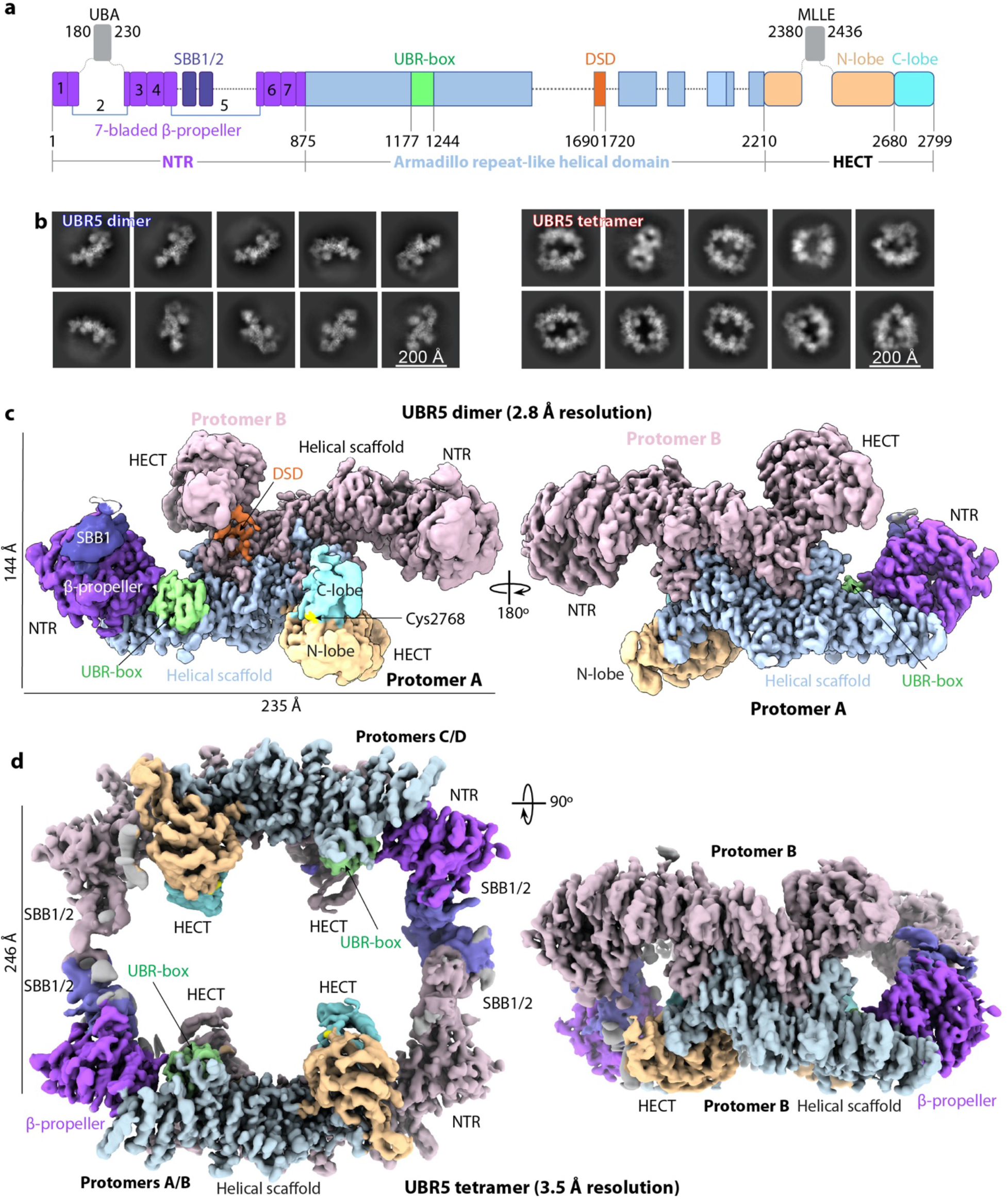
Cryo-EM of the UBR5 dimer and tetramer. **a**, Domain architecture of the human E3 ligase UBR5. Unresolved regions in the EM map are shown as dash lines. The disordered UBA and MLLE domains are shown as gray squares. **b**, Selected 2D class averages of the dimer (left) and tetramer (right). **c-d**, Cryo-EM 3D maps of the dimer (c) and tetramer (d). Domains are colored as in (a).

## RESULTS

### UBR5 assembles a dimer and a tetramer as revealed by cryo-EM

We recombinantly expressed human UBR5 with an N-terminal FLAG tag in insect cells and purified the protein via affinity and size exclusion chromatography. Purified UBR5 eluted from the gel filtration column in two broad peaks at volumes corresponding to a dimer (620 kDa) and a tetramer (1.2 MDa) (**Supplementary Figure 1a-b**). PEPCK1 was previously shown to be a UBR5 substrate in vivo (Jiang et al., 2011; Shen et al., 2018). We demonstrated that the purified UBR5 was enzymatically active as it ubiquitylated PEPCK1 in vitro in the presence of Ub, E1, and E2D2, while the catalytically inactive UBR5 mutant protein (C2768S) was unable to do so (**Supplementary Figure 1c**). Cryo-EM and 2D image classification revealed a predominant dimer species and a subset of particles that are dimer-of-dimers tetramers (**Supplementary Figure 1d, Fig. 1b**). We derived a cryo-EM 3D map of the dimer at an average resolution of 2.8 Å in C1 symmetry with an overall dimension of 108 Å × 144 Å × 235 Å (**Fig. 1c**, **Supplementary Figures 2-3, Supplementary Table 1**). Resolution increased to 2.66 Å when 2-fold symmetry was applied. However, the two ends of the dimer map, corresponding to the N-terminal region (NTR) and the HECT region, are of lower resolution. Therefore, we performed focused refinement on the two regions separately, and improved the map quality in these regions, leading to a high-resolution C2 symmetric composite EM map of the dimer (**Supplementary Figures 2, 4**). Next, we combined an untilted and a 30° tilted dataset to obtain a composite EM map of the UBR5 tetramer at 3.5 Å average resolution (**Supplementary Figures 5-6**). The 3D map was determined in the C1 symmetry, because application of the C2 or D2 symmetry led to a loss of densities in the HECT and NTR, indicative of mobility in these regions. The tetramer is assembled by interaction between the NTR regions of the two dimers, leading to more density but not increased resolution of the junction region. The tetramer structure has an overall dimension of 221 Å × 238 Å × 126 Å, with the four catalytic HECT domains all facing inside the large central cavity (**Fig. 1d**).

### The UBR5 dimer structure

The high-quality EM map of the UBR5 dimer allowed us to build an atomic model for most regions of UBR5, except for the UBA, SBB2, and MLLE domains that are flexibly linked to the core of the dimer complex (**Fig. 2a**, **Supplementary Figure 7, Supplementary Video 1**). Two UBR5 subunits assemble head-to-tail via their respective helical scaffold to form an extended dimer interface of 7936 Å^2^. The crystal structures of isolated UBA and MLLE domains are known (Kozlov et al., 2007; Muñoz-Escobar et al., 2015). UBA is flexible in the UBR5 dimer because it is tethered to the NTR by a preceding 100-residue linker and a following 120-residue linker. Contrary to a previous bioinformatic analysis, MLLE does not precede the HECT, instead it is inserted into the N-lobe of the HECT domain and is flexibly connected by a 63-residue-long linker on each side. Strikingly, the NTR of one subunit is close to the HECT domain of the other subunit, forming a gap that we speculate could bind an E2–Ub. For this reason, we tentatively term the gap an “intermolecular E2–Ub jaw”. Individual UBR5 molecules adopt a crescent shape with a bridging helical scaffold (residues 876-2209) interrupted by the well-ordered E3 substrate N-terminal recognition UBR-box (1177-1244). The NTR (1-875) and the C-terminal HECT domain (2210-2799) straddles the two ends of the rigid helical scaffold platform, enclosing a space for E3 substrate binding.

**Fig. 2.**
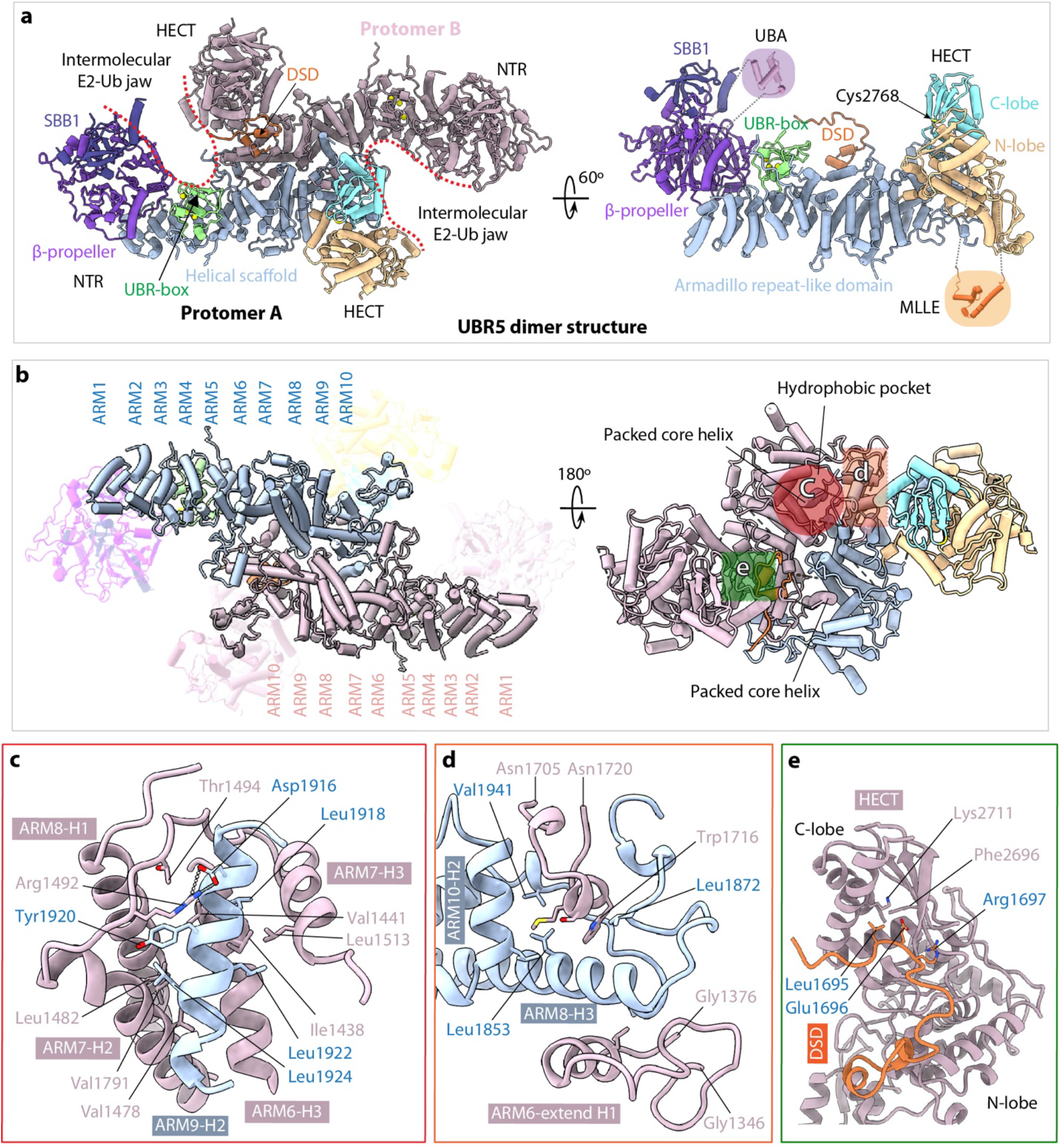
The UBR5 dimer structure. **a**, The dimer structure in cartoon viewed from top along the 2-fold symmetry axis (left) and a monomer in side view (right). Th structures are colored as in Fig. 1a. The crystal structures of the UBA (PDB ID 2QHO) and MLLE (PDB ID 3NTW) are shown in shadowed cartoons for illustrative purpose only; they are invisible in the EM map. **b**, Top and bottom views of the middle Armadillo-like helical scaffold that primarily mediates UBR5 dimerization. The three major interacting regions are marked by three colored shapes in the right panel. **c**, Close-up view of the hydrophobic interface region marked by the red circle in (b). Residues involved in dimerization such as the salt bridge between Arg1492 and Asp1916 are shown as sticks. **d**, Close-up view of the interface region marked by the orange square in (b). This region contains both hydrophobic and H-bonding interactions. **e**, Close-up view of the region marked by green square in (b), which involves the domain-swapped dimerization (DSD) motif.

The helical scaffold is composed of 10 Armadillo-like (ARM1-10) repeats (**Fig. 2b**). The second half of the helical scaffold, ARM5-10, is near the 2-fold symmetry axis and is primarily responsible for the dimer interface. The two symmetry-related ARM5-10 motifs interact with each other hydrophobically. Specifically, the ARM9 interface helix 2 (ARM9-H2) of one subunit inserts into an extended hydrophobic pocket formed by ARM6-8 of the partner subunit; this interface is further stabilized by a salt bridge between Arg1492 of one subunit and Asp1916 of the other (**Fig. 2b-c**). The peripheral dimer interface is also largely hydrophobic and is formed by the ARM8 H3 helix inserting into a space in the partner subunit between the ARM6 extended H1 helix and an extended loop (1705-1720) (**Fig. 2d**). Interestingly, the dimer interface is further stabilized by a domain-swapped dimerization loop (DSD loop, aa 1690-1700) interacting with both N- and C-lobes of the HECT domain (**Fig. 2e**). The presence of the DSD loop in the catalytic domain likely has functions beyond dimerization, because E3 ligase activity involves the relative movement of the N- and the C-lobes, as described below.

### The N-terminal region (NTR)

The NTR of a UBR5 dimer is located above ARM1-4 that are not involved in the helical scaffold dimerization (**Fig. 3a-b**). The NTR is composed of SBB1 and β-propeller with SBB1 sitting above the β-propeller. The SBB1 contains five β-strands with an SH3-like fold. The SH3 fold is typically involved in protein-protein interaction or nucleic acid binding (Youkharibache et al., 2019) or in substrate degron recognition (Sherpa et al., 2022). The UBR5 SBB1 lines the intermolecular E2-Ub jaw and could be involved in substrate binding. The β-propeller domain is seven-bladed with a central pore, with each blade containing four β-strands (**Fig. 3b**). Blade 1 is composed of the β-strands from both ends of the NTR (**Fig. 1a, Supplementary Figure 7**) and contains an extra α-helix that connects the β-propeller domain with the middle helical scaffold. Blades 2 to 7 are arranged clockwise to complete the β-propeller fold with blade 1. Blade 2 features two extended loops that link the unresolved, mobile UBA domain. SBB1-2 are inserted into blade 5. Finally, blade 7 interacts with the UBR-box (**Fig. 3a**). β-propeller is also often involved in protein-protein interaction (Chen et al., 2011), and several multi-protein E3s including KLHDC1-3, KLHDC10, and Gid11 recruit either C-degrons or N-degrons to the central pore of their respective β-propellers (Sherpa et al., 2022). However, the central pore of the UBR5 β-propeller is partially covered by SBB1 on top. Because the C-terminal catalytic HECT and MLLE directly interact with UBR5 E3 substrates (Jiang et al., 2015; Shen et al., 2018; Yoshida et al., 2006), it will be interesting to investigate if the NTR (SBB1/2 and β-propeller) may interact with other protein partners, given that UBR5 functions well beyond the E3 ligase (Liao et al., 2017; Song et al., 2020; Xiang et al., 2022).

**Figure 3.**
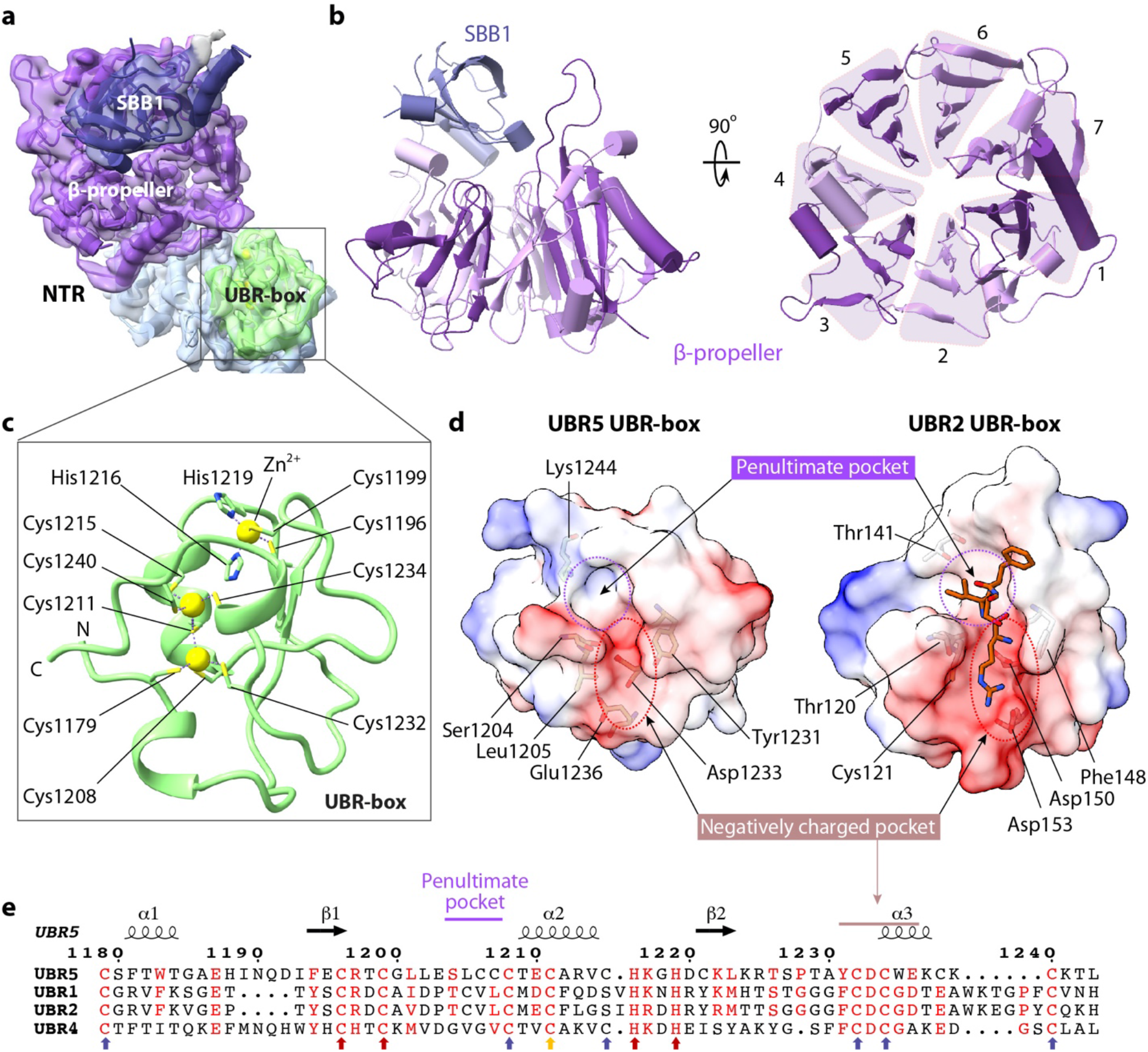
The NTR and UBR-box structures. **a**, Focus-refined EM map of the NTR and UBR-box in transparent surface view superimposed with atomic model in cartoons and colored as in Fig. 1b. **b**, Left: Side view of the β-propeller and the small β-barrel 1 (SBB1) in the NTR. Right: Top view of the seven-blades β-propeller. **c**, Structure of the UBR-box with two zinc fingers coordinating three zinc ions (yellow spheres). The coordinating cysteine and histidine residues are in sticks. **d**, Electrostatic surface views of the UBR5 UBR-box (left) and the UBR2 UBR-box bound to an N-degron peptide shown in orange sticks (right, PDB ID 3NY3). The two N-degron binding subsites are marked by dashed red and purple circles, respectively. **e**, Sequence alignment of the UBR boxes of human UBR5, UBR1, UBR2, and UBR4. The conserved Cys and His residues coordinating the first Zn^2+^ are indicated by red arrows. The six Cys that coordinate the remaining two Zn^2+^ are indicated by blue arrows. Cys1211 participates in coordination of two zincs and is indicated by a yellow arrow.

### The UBR-box contains a conserved N-degron recognition pocket

So far, all known structures of the UBR-box are from the RING E3 ligases (Choi et al., 2010; Matta-Camacho et al., 2010; Munoz-Escobar et al., 2017; Pan et al., 2021). To our knowledge, the UBR5 structure provides the first HECT-family UBR box structure. The UBR5 UBR-box coordinates three zinc ions with two zinc fingers having little regular secondary structure (**Fig. 3c**), which is similar to other UBR-box structures. The first zinc finger is a typical Cys_2_His_2_ type, with Cys1196, Cys1199, His1216, and His1219 coordinating the top zinc ion. The second zinc finger contains the middle and the bottom zinc ions. The middle zinc is coordinated by Cys1179, Cys1208, Cys1232, and Cys1211, and the bottom zinc is coordinated by Cys1211, Cys1215, Cys1234, and Cys1240. Therefore, Cys1211 participates in the coordination of two zinc ions. This Zn_2_/Cys_7_ coordination differs from the well-known binuclear zinc clusters, such as the Zn_2_/Cys_6_ pattern observed in the yeast Gal4 protein or the Zn_2_/Cys_6_-His pattern observed in the RING UBR boxes (Sriram and Kwon, 2010) (**Supplementary Figure 8**).

The UBR5 UBR-box contains a negatively charged pocket for recognizing the N-terminal residue and a hydrophobic pocket for the second residue (**Fig. 3d**), consistent with the knowledge that UBR5 targets type 1 N-degrons in which the first amino acid is positively charged Arg, Lys, or His (Sherpa et al., 2022; Sriram and Kwon, 2010). The two amino acid binding pockets in a peptide bound UBR2 UBR-box structure are larger than those of the UBR5 UBR-box (**Fig. 3d**), due to the presence of residues with longer side chains in UBR5. The negatively charged pocket is well conserved among E3s with the N-degron related UBR box (**Fig. 3e**). The penultimate hydrophobic pocket is also conserved, but to a lesser extent.

### The UBR5 tetramer is assembled via SBB2-SBB2 interaction between two dimers

The 3D EM map of the UBR5 tetramer readily fits two UBR5 dimers that fuse into a cage like structure (**Fig. 4a**). Thus, the tetrameric architecture places the four UBR-boxes and four HECT domains all facing the inside of the large central chamber. This organization suggests that the UBR5 tetramer may encapsulate a substrate and add multiple Ub to the substrate simultaneously, like the GID E3 chelator ^17^. The primary interfaces between the two dimers are clearly located in the SBB2 region, which is not resolved in the UBR5 dimer structure. We used the AlphaFold-Multimer to predict the SBB1-2 structure and the dimerization interface (Jumper et al., 2021). The predicted interface between two SBB1-2 structures is of high confidence and consistent among the top five predicted structures (**Supplementary Figure 9**). Importantly, the predicted SBB1-2 dimer structure fits well with the EM density at the dimer-dimer interface (**Fig. 4b**). Therefore, the tetramer is mediated by intermolecular SBB2-SBB2 interaction. The interface is stabilized by hydrophobic interactions involving Pro701, Leu705, and Leu710, as well as H-bonds between Glu715 and Tyr677 and between Asp674 and the backbone nitrogen atom of Leu710 (**Fig. 4c**). The tetramer interface is ~800 Å^2^ per junction, much smaller than the dimer interface, suggesting that the tetramer is less stable than the dimer in solution.

**Figure 4.**
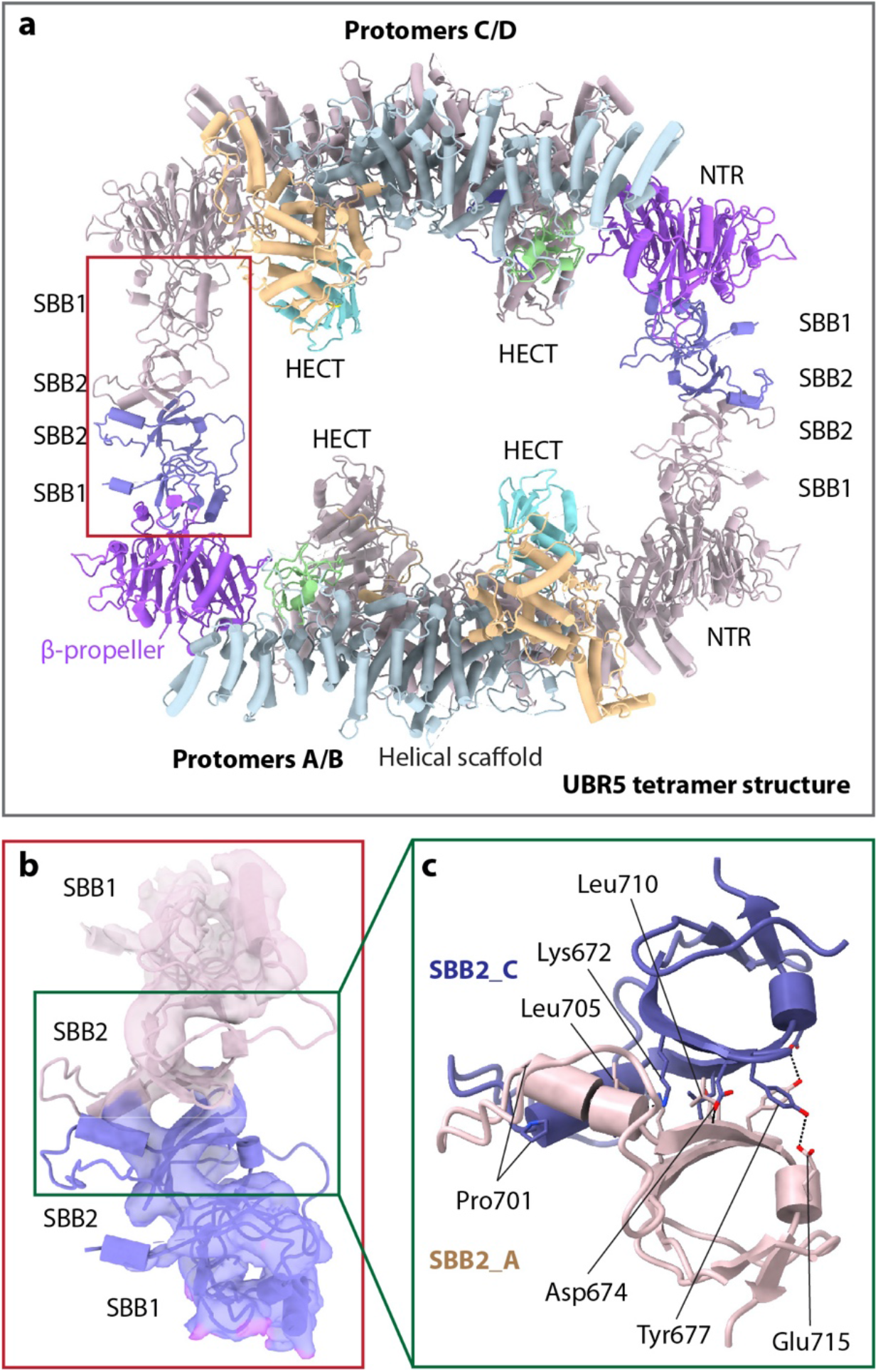
The UBR5 tetramer structure. **a**, Atomic model of the tetramer UBR5 in cartoon view. Two UBR5 chains (A and C) are colored as in Fig. 1c, and the two remaining chains in salmon. The red rectangle marks the tetramerization interface between two dimers that is mediated by the SBB2-SBB2 interaction. **b**, Close-up view of the red rectangle region in panel a showing the EM density of SBB1/2 of protomers A and C in transparent surface superimposed with atomic model in cartoons. **c**, Close-up view of the interface in the green box in panel b showing the SBB2 residues involved in tetramerization as predicted by AlphaFold-multimer.

### The UBR5 dimer undergoes large conformational changes around the E2-Ub jaw

The N-lobe and C-lobe of the HECT domain are connected by a flexible linker. In the truncated homologous HECT domains, these two lobes were previously shown to arrange either in an L or an inverted T conformations based on the orientation of the C-lobe with respect to N-lobe (Kamadurai et al., 2013; Kamadurai et al., 2009; Lorenz, 2018; Maspero et al., 2011; Maspero et al., 2013; Rotin and Kumar, 2009; Singh et al., 2019). The UBR5 HECT is in the L-conformation (**Fig. 5a**), the Ub–E2 bound NEDD4L HECT is in the inverted-T configuration with the C-lobe sitting in the middle of the long axis of the N-lobe (**Fig. 5b**) (Kamadurai et al., 2009; Rotin and Kumar, 2009). When this structure is superimposed with UBR5 HECT, the E2-linked Ub is in the backside of UBR5 C-lobe (**Fig. 5c**). The UBR5 C-lobe needs to rotate ~130° to align with the C-lobe of NEDD4L HECT that interacted with the Ub–E2. We asked if the UBR5 dimer indeed undergoes large conformational changes as suggested above. We performed a 3-dimensional variability analysis (3DVA) (Punjani and Fleet, 2021). We found that the apo UBR5 dimer in solution fluctuates among a series of conformations **(Supplementary Video 2)**, with the NTR and HECT undergoing the most significant motions, including the lateral expansion of the NTR and an up to 55° rotation of the HECT domain (**Fig. 5d**). These intrinsic motions are likely related to and perhaps enable the above-described conformational changes based on structural comparison (**Fig. 5c**). When the structure of the Ub–E2 bound NEDD4L HECT is superimposed with the UBR5 HECT in the UBR5 dimer, the Ub–E2 fits well inside the E2–Ub jaw (**Fig. 5e**), indicating that the UBR5 intermolecular jaw is most likely involved in the Ub transferring from E2–Ub to UBR5 (**Fig. 5f**).

**Figure 5.**
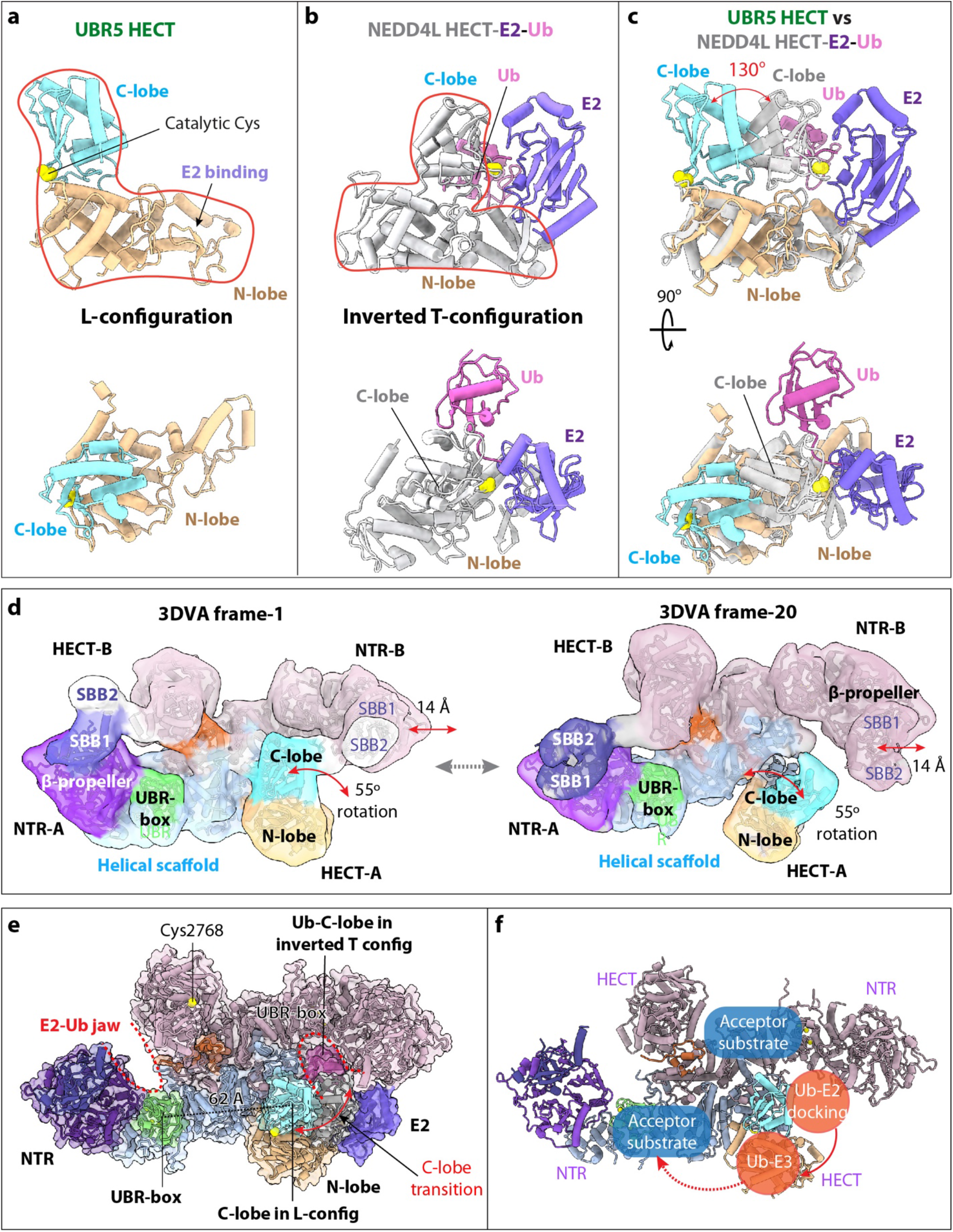
UBR5 undergoes large conformational changes around the E2-Ub jaw. **a**, UBR5 HECT domain in the L-conformation in two orthogonal views. **b**, The NEDD4L-HECT-E2-Ub structure (PDB ID 3JWO) in the same view. **c**, Superimposition of the UBR5-HECT (this study) and NEDD4L-HECT-E2-Ub (PDB ID 3JWO) structures. The UBR5 HECT N-lobe is poised to bind E2, but the C-lobe needs to rotate 130° to reach the C-lobe position of the NEDD4L-HECT for transthiolation reaction. **d**, The two most distinct conformations of the UBR5 dimer as determined by 3DVA, showing a 14 Å lateral movement of the NTR and a 55° rotation of the HECT. SBB2 above SBB1 was observed in this lower resolution variability analysis, but missing in the 2.8 Å 3D map, indicating its high mobility in the dimer. **e**, Ub-E2 docked in the right intermolecular jaw of the UBR5 dimer. The distance between the C-lobe and UBR-box is 62 Å. **f**, Possible substrate ubiquitylation pathway. The curved red arrow indicates that E3 Ub transthiolation reaction occurs in the intermolecular jaw. The dashed red arrow indicates the Ub transfer route for substrate ubiquitylation.

### Residues lining the E2-Ub jaw are important for UBR5 ligase function

Motions of the jaw-forming NTR and HECT point to their function in the initial Ub-loaded E2 recruitment for transthiolation as well as in the following E2 release and Ub-linked HECT reorientation for substrate ubiquitylation. To gain insights into this process, we first identified jaw-lining residues that potentially interact with Ub or E2. Superimposition of our UBR5 HECT with published Ub bound or Ub–E2 bound HECT structures showed that UBR5 HECT N-lobe likely binds E2 (UbcH5B) via hydrophobic interactions involving Met2575 and Tyr2576, and UBR5 C-lobe binds Ub via hydrophobic interactions involving Phe2732, Leu2789, and Ala2790 (**Fig. 6a**). The published UBR5 UBA-Ub complex structure has revealed the hydrophobic interactions involving UBR5 Leu8, Val196, and Leu224 (Kozlov et al., 2007). Therefore, we made five single mutations lining the “jaw”, V196K, L224K, Y2576A, F2732A, and A2790W, plus the active site mutation C2768S, purified the mutant UBR5 proteins, and performed the E2 discharge assay. In this assay, we labeled Ub with a fluorescent dye, loaded Ub by E1 onto a E2 (E2D2), then added purified WT and mutant UBR5 proteins, and monitored the transthiolation reaction in which Ub was transferred from E2 to UBR5. We found the WT UBR5 had robust Ub discharge activity from E2 and the catalytically inactive C2768S mutant protein had no activity (**Fig. 6b-c**). The A2790W and F2732A substitutions at the catalytic C-lobe interface with Ub had little activity. Single mutation Y2576A in the E2-interacting HECT N-lobe reduced but did not abolish the UBR5 activity. The UBA domain is inserted in the NTR β-propeller. We found that mutating the two Ub-interacting residues (V196K and L224K) reduced the activity. This result is consistent with the previous finding that substitution of V196 and L224 disrupted interaction between UBA and Ub (Kozlov et al., 2007). We further generated an NTR-deletion UBR5 (Δ1-875) and found that the truncation compromised but did not abolish the activity. These results support our proposal that the intermolecular jaw is important for UBR5 function.

**Figure 6.**
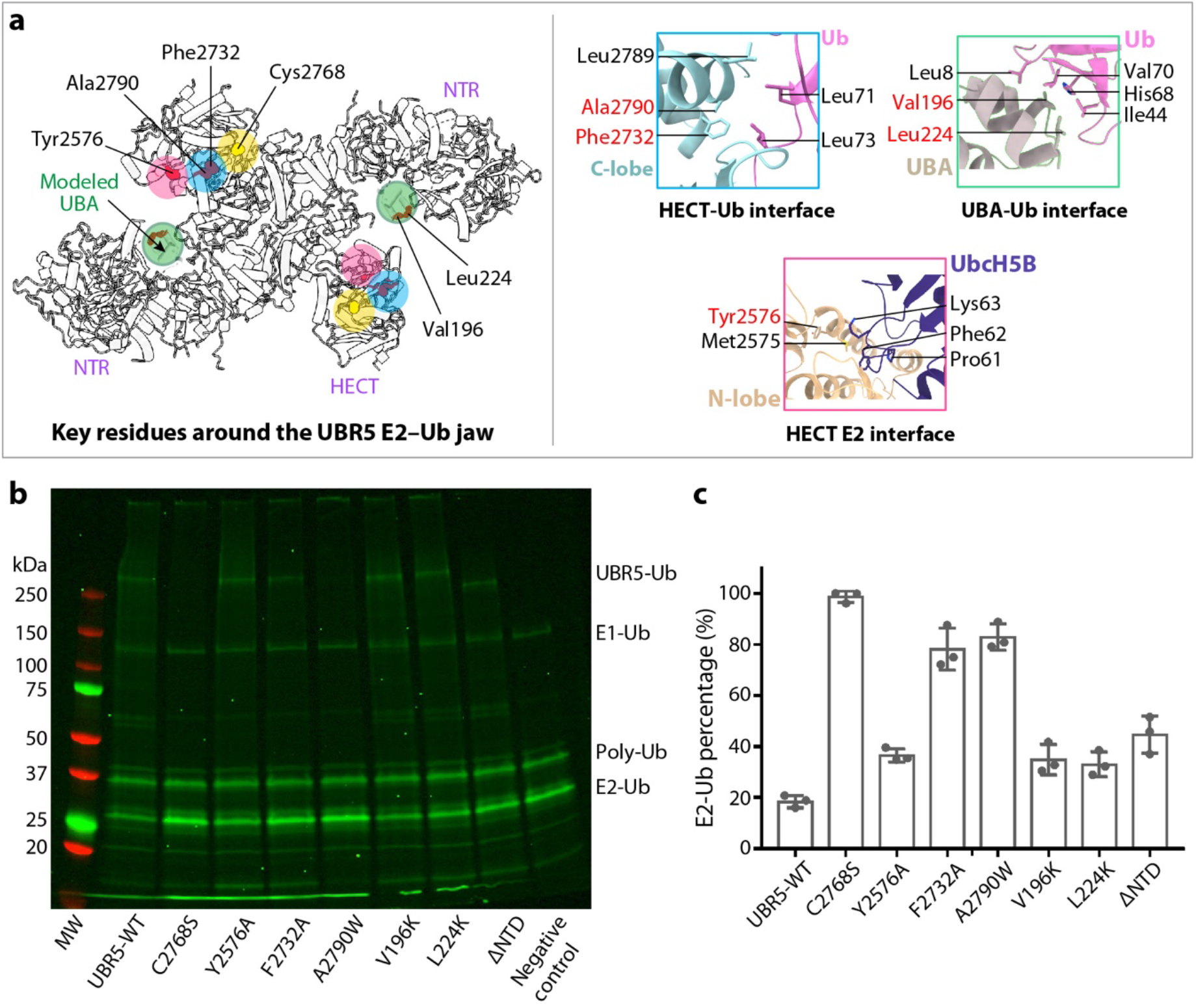
Ub transfer assay from E2-Ub to UBR5 by WT and mutations around the E2-Ub jaw. **a**, Key residues involved in E2 and Ub binding are displayed as red spheres, and their locations are highlighted by colored circles. The catalytic cysteine is in yellow. The right panels show the catalytic pocket and three predicted interfaces between HECT C-lobe and Ub, between HECT N-lobe and Ub, and between UBA and Ub, based on alignment with the isolated HECT–Ub structures shown in Fig. 5a. The UBA location is based on the published isolated UBR5 UBA–Ub complex structure (PDB ID 2QHO). **b**, In-gel fluorescence of the E2 discharge assay by purified WT and seven mutant UBR5 proteins. **c**, Quantification of the E2 discharge assay results. All values represent means ±SD obtained from three independent experiments. ΔNTD is a truncated UBR5 removing residues 1-875.

To further investigate the importance of the jaw-lining residues in substrate ubiquitylation in vivo, we co-transferred plasmids expressing UBR5 mutants with a UBR5-targeting shRNA (shUBR5) into the MDA-MB-231 cells and analyzed the CDC73 protein levels by Western blotting (**Fig. 7a**). The tumor suppressor CDC73 is a component of RNA polymerase II-associated factor 1 complex and has been demonstrated a UBR5 substrate ^28^. As a control, we first confirmed that shUBR5 drastically suppressed the UBR5 expression level with a concomitant accumulation of CDC73. The normalized expression levels of most UBR5 mutants were similar to that of the vector control, except for the A2790W, V196K, and C2768S mutants that were somewhat higher (**Fig. 7b**). However, the CDC73 levels in cells expressing UBR5 mutants were much higher than that of the empty vector control, except for Y2576A (**Fig. 7c**). Therefore, most of the E2 and Ub interacting residues had diminished E3 activity towards CDC73. And this in vivo result is consistent with the in vitro E2 Ub discharge assay (**Fig. 6c**). Taken together, our in vitro and in vivo data demonstrate that the E2-Ub jaw plays an important role in the UBR5 function.

**Figure 7.**
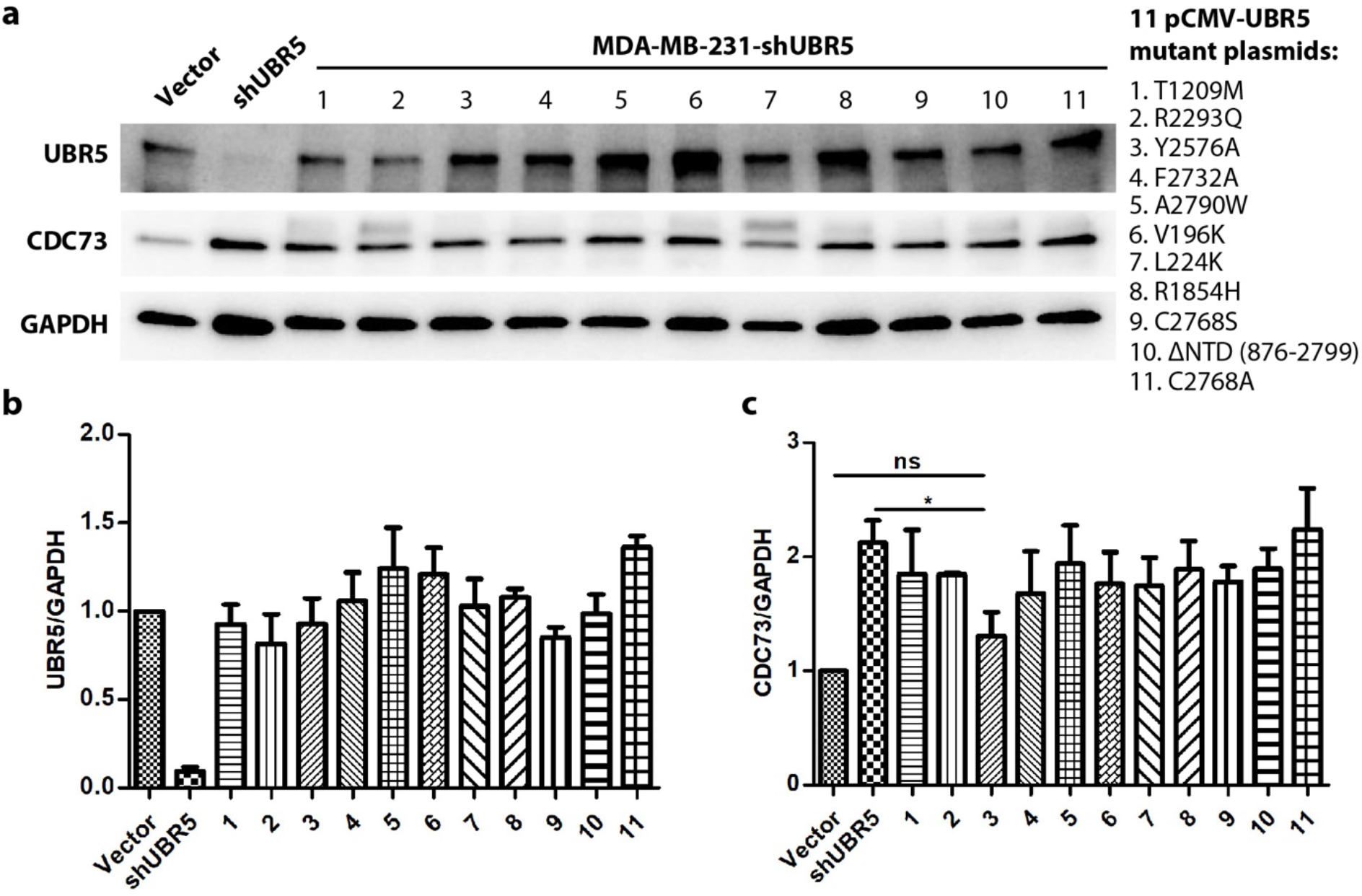
*In vivo* ubiquitylation assay on CDC73 by the WT and mutant UBR5. **a**, The protein level of UBR5 (top), the substrate protein CDC73 (middle), and the control GAPDH (bottom) in the MDA-MB-231 cells were detected by Western blotting. **b**, Quantification of the in *vivo* protein levels of UBR5 from (**a**). **c**, Quantification of the in *vivo* protein levels of CDC73 from (**a**). Data from three independent experiments.

## DISCUSSION

Our cryo-EM analysis has shown that a UBR5 monomer adopts a crescent shape comprising ten ARM repeats forming the central helical scaffold, an NTR containing multiple potential protein-protein interaction modules, and a HECT domain that ubiquitylates target substrates (Lorenz, 2018). Unexpectedly, we found that UBR5 oligomerizes into a dimer and a tetramer. While UBR5 dimerization is mediated by the central helical scaffold, leaving both NTR and HECT domains unconstrained by the dimerization interface, UBR5 tetramerization is mediated by intermolecular SBB2-SBB2 interaction between two UBR5 dimers. Because the SBB2 interface is small relative to the dimer surface, the tetramer is likely less stable than the dimer. It is possible that the tetramer interface may open up to either become a dimer or recruit more dimers to assemble a larger cage, as found in the multiprotein GID E3 complex ^17^.

Previous studies have suggested that Ub transfer from E2 to E3 HECT occurs when the HECT is in the inverted T conformation and Ub transfer from E3 HECT to substate occurs when the HECT is in the L-conformation (Kamadurai et al., 2013; Kamadurai et al., 2009; Lorenz, 2018). Our 3DVA analysis of UBR5 dimer shows the C-lobe of HECT undergoes larger scale rotation in the absence of a E2–Ub (**Fig. 4a**). We suggest such intrinsic dynamism may account for the UBR5’s ability to transition from the L to inverted T conformation during catalysis. Our structural and functional studies suggest that the first step of the Ub transthiolation reaction between the Ub–E2 and UBR5 occurs inside the intermolecular jaw formed between the NTR of one subunit and the HECT of the partner subunit in the inverted T conformation. Upon accepting Ub from E2 to form the Ub-UBR5 intermediate, the catalytic C-lobe likely reverts to the observed L conformation. The next step of ubiquitylation reaction occurs between the E3-bound Ub and a protein substrate. The mechanism of the second Ub transfer is complicated by UBR5 oligomerization. In the case of the UBR5 dimer, the E3-bound Ub is likely transferred to a substrate bound to the same UBR5 subunit (**Fig. 5e**).

The E3 activity can be regulated by multiple factors, including post-translational modifications, intermolecular and intramolecular interactions, and accessory proteins or adaptors (Gallagher et al., 2006; Ogunjimi et al., 2005; Sander et al., 2017; Shah and Kumar, 2021; Wiesner et al., 2007). In UBR5, the enzyme’s ability to form dimer and tetramer is notable. Of particular interest is the domain-swapped dimerization (DSD) motif projecting from one subunit into the HECT domain of the partner subunit. This motif wedges between the N-lobe and C-lobe at the back of the partner HECT domain. Because the C-lobe is expected to undergo large conformational changes during catalysis, the DSD motif likely regulates the UBR5 activity. In the UBR5 tetramer, the DSD motif could extend into the E2–Ub jaw to potentially influence the tetramer interface, thereby regulating the tetramer assembly (**Fig. 1d**). Further studies are required to determine how oligomerization affects the UBR5 activity, and whether there are distinct biological functions or substrates between the dimer and the tetramer.

## MATERIALS AND METHODS

pET28-mE1 was a gift from Jorge Eduardo Azevedo (Addgene plasmid # 32534; http://n2t.net/addgene:32534; RRID: Addgene 32534). The codon-optimized human UBR5 with an N-terminal Flag tag, human His6-E2D2, and human CDC73 sequences with an N-terminal Flag tag were synthesized and sequences confirmed by GenScript. The human ubiquitin sequence with an inserted N-terminal cysteine residue was synthesized by Eurofins. The PEPCK1 expression plasmid was a gift from Dr. Lewis Cantley at Harvard Medical School.

### Protein expression and purification

The human UBR5 sequence with an N-terminal Flag tag was cloned into pFastBac with a polyhedrin promoter. The UBR5 mutants were generated by PCR based mutagenesis. The NTR deletion of UBR5 (876-2799 aa) was cloned into the pFastBac vector using the same restriction sites as the UBR5 wild type. UBR5 and UBR5 mutants were expressed in Sf9 insect cells (Novagen) using the Bac-to-Bac baculovirus expression system (ThermoFisher). Cell cultures were grown in ESF 921 serum-free medium (Expression Systems) to a density of 2.5 × 10^6^ cells/mL and then infected with the UBR5 baculovirus (3%-5% v/v) for 48-60 h post-transfection. We found the use of a higher UBR5 baculovirus infection ratio and the longer time after transfection leads to a higher population of UBR5 tetramer. The cells were collected, flash frozen, and stored at –80 °C for further usage. Cells were lysed by sonication in lysis buffer (25 mM HEPES, PH7.5, 200 mM NaCl, 10% glycerol, 1 tablet EDTA-free protease inhibitors cocktail), centrifugated in a Ti-45 rotor at 40,000x rpm for 1 h, and incubated with FLAG-antibody-coated beads at 4 °C for 2-3 h. Beads were washed with 50 column volume (CV) lysis buffer, and the proteins were eluted with 5 CV of lysis buffer with 0.2 mg/mL 3 × FLAG peptides. The proteins were concentrated using centrifugal concentrators (Amicon, 100 kDa) and polished by size exclusion chromatography (SEC, Superose 6 Increase, GE Healthcare) in buffer containing 25 mM HEPES, PH 7.5, 100 mM NaCl, 0.5 mM TCEP. Fraction peaks corresponding to the dimer and tetramer were pooled separately for cryo-EM analysis.

The mouse E1 and human E2D2 were cloned into the pET22b vectors. Ub with an added N-terminal cysteine residue was cloned into a pET28a vector to make Ub with an N-terminal 6xHis-tag and a thrombin cleavage site. The human PEPCK1 was cloned into the pET28a vector with an N-terminal His tag. These plasmids were individually transformed into *E. coli* BL21 (DE3) cells (Life Technologies). All recombinant strains were grown in 2 L LB medium at 37°C. When the cell density reached an OD_600_ value of 0.8, 0.2 mM isopropyl-β-D-thiogalactopyranoside (IPTG) was added, and the culture was continued for 12 h at 16°C. We then collected the cells, resuspended the cells in buffer A (20 mM HEPES pH 7.6, 150 mM NaCl, and 10% glycerol), and lysed the cells with a homogenizer (SPX Corporation). The lysate was centrifuged at 20,000x g for 40 min, and the supernatant was collected and loaded into a 5-ml Ni-NTA column (Cytiva). The proteins were eluted using buffer A plus 300 mM imidazole. The E1, E2, and PEPCK1 proteins were further purified by a Superdex 200 column in buffer A. The N-terminal 6xHis-tag on Ub was cleaved by incubating with thrombin at 4°C overnight. The tag cleaved Ub was further purified by size exclusion chromatography through a Superdex 75 column (GE Healthcare) in buffer A.

### Cryo-EM sample preparation and data collection

The holey carbon grids (Quantifoil Au R2/1, 300 gold mesh) were glow discharged in the Ar/O2 mixture for 30 s using a Gatan 950 Solarus plasma cleaning system with a power of 15 W. Aliquots of 3 μl of purified UBR5 solution at a concentration of 0.7 mg/ml were placed on the freshly treated EM grids. The grids were blotted with qualitative cellulose filter paper (TED PELLA, INC) for 3 s after each sample application with the blotting force set to 3 and flash-frozen in liquid ethane using an FEI Vitrobot Mark IV. Temperature and relative humidity were maintained at 6°C and 100%, respectively. We prepared cryo-EM grids for both the dimer and tetramer elution peaks then loaded the grids into an FEI Titan Krios electron microscope operated at a high tension of 300 kV. Cryo-electron micrographs were collected automatically with the SerialEM program (Mastronarde, 2018) at a nominal magnification of 105,000 X in a K3 summit direct electron detector (Gatan) in multi-hole mode, with the objective lens defocus being varied between –1.0 to –2.0 μm. The K3 camera was operated in the super-resolution counting mode. During a 1.5-s exposure time, a total of 75 frames were recorded with a total dose of 65 e^-^/Å^2^. The calibrated physical pixel size was 0.828 Å at the specimen level for all digital micrographs. All datasets were collected with the sample stage tilt angle set to 0°, except for one dataset of the UBR5 tetramer that was recorded with a tilt angle of 30° to alleviate the preferred orientation problem of the tetramer.

### Image processing

For the untilted dataset, a total of 13,090 raw movie micrographs were collected and motion-corrected using the program MotionCorr 2.0 (Zheng et al., 2017). Then the micrographs were imported into cryoSPARC (version 3.2.0) (Punjani et al., 2017) to perform the patch-based contrast transfer function (CTF) estimation and correction. A total of 12,471 micrographs with CTF signals extending to 4.1 Å were retained in further processing. We first used the blob particle picking (70–300 Å diameter) to generate 2D templates for subsequent automatic particle picking. In total, 4,262,102 particles were automatically picked and extracted. Two rounds of 2D classifications were then performed on particle images with 4x binning and particles in the classes with clear structural features were selected. A total of 1,931,379 particles was used to calculate five starting 3D models. One major class was chosen to perform a new round of 3D classification to remove bad particles. Two major 3D classes were combined to preform further 3D refinement and post-processing, resulting in the 2.80-Å 3D density map with 844,403 particles in C1 symmetry. Application of 2-fold symmetry during 3D reconstruction and refinement led to the final 3D map at an overall resolution of 2.66 Å. The map resolution was estimated by the gold-standard Fourier shell correlation at a correlation cut-off value of 0.143 (Scheres and Chen, 2012). We further performed focused refinement on the NTR and the HECT region separately. The focus refined NTR and HECT maps had an average resolution at 2.94 Å and 3.01 Å, respectively. Finally, we combined the focus-refined maps with the original C2 symmetric map to generate a composite EM map for the UBR5 dimer.

Because the NTR and the C-terminal HECT region were of lower resolution, implicating dynamics in these regions. We next used the 3D variability analysis (3DVA) program (Punjani and Fleet, 2021) to investigate the conformational heterogeneity in the UBR5 dimer. The final dataset with 844,403 particle images were down sampled by a factor of 4 and subjected to non-uniform refinement in C1 symmetry to generate an appropriate mask. 3DVA was carried out to solve three principal components with the filter resolution set to 8 Å; and other parameters used their default values. The distribution of reaction coordinates across particles in the principal components were smooth, confirming the compositional homogeneity of the UBR5 dataset and the conformational flexibility of the NTR and HECT domain.

The 30° tilted dataset of the tetramer peak sample contained a total of 17,373 raw movie micrographs. The micrographs were motion-corrected using the program MotionCorr 2.0(Zheng et al., 2017) and combined with the untitled micrographs imported into cryoSPARC (version 3.2.0) (Punjani et al., 2017) to perform the patch-based contrast transfer function (CTF) estimation and correction. A total of 30,467 micrographs with CTF signals extending to 4.0 Å were retained for further process. We also generated 2D templates for the tetramer particles from the untilted dataset and used template picking (290 Å diameter) to select an initial dataset of 1,793,574 tetramer particles. Several rounds of 2D classifications were then performed on 4x binned particle images. Particles in the 2D classes with clear structural features were retained, resulting in a selected dataset of 1,027,335 particles, which was used to calculate three starting 3D models. We performed heterogenous refinement on the particle images belonging to the best start model and obtained 3 additional maps; particles belonging to the two good maps were then combined (401,468 particles) for further refinement and post-processing, resulting in the 3.5-Å 3D density map in C1 symmetry. Application of 2-fold symmetry or 4-fold symmetry during 3D reconstruction and refinement led to a high-resolution 3D map but with reduced densities in certain regions. The resolution of the C1 map was estimated by the gold-standard Fourier shell correlation at a correlation cut-off value of 0.143 (Scheres and Chen, 2012). We further performed focused refinements on two UBR5 dimers and the dimer-dimer interface regions separately. The focus refined UBR5 dimer maps and the interface region map had an average resolution of 3.4 Å. We finally combined these partial maps with the original C1 map to generate a composite 3D map for the UBR5 tetramer.

### Model building, refinement, and validation

We first used the AlphaFold2 server installed in a local workstation to obtain a predicted UBR5 atomic model (Jumper et al., 2021). The high-confidence regions, including the N-terminal small β-barrel 1 (SBB1) and β-propeller, the UBR box, the helical repeat, and the C-terminal HECT domain, were docked individually into the cryo-EM maps using UCSF Chimera (Pettersen et al., 2004). We then manually adjusted each domain and built missing residues in our high-resolution cryo-EM map in Coot (Emsley and Cowtan, 2004). We also referenced to the high-resolution (2.66 Å) 2-fold symmetric composite 3D map to build several regions with weaker densities. The completed model was subjected to several iterations of real-space refinement in PHENIX (Adams et al., 2010) and manual adjustment in Coot (Emsley and Cowtan, 2004). For modeling the UBR5 tetramer map, two dimer models were readily fitted into the composite EM map. But the dimer-dimer interface region had a lower resolution that was insufficient for manual model building. We used the AlphaFold-Multimer server to predict a dimer structure of the SBB1-SBB2 sequence. All predicted dimer structures had a high confidence value over 95% with a consistent SBB2-SBB2 interface. The predicted SBB2-SBB2 complex structure was fitted well in the dimer-dimer interface region of the UBR5 tetramer map. Finally, all atomic models were validated using MolProbity (Williams et al., 2018). The original EM maps in both C1 and C2 symmetry and the maps sharpened by deepEMhancer were used to refine the atomic models (Sanchez-Garcia et al., 2021). The statistics of the model refinement are shown in **Supplementary Table 1**. Structural figures were prepared in UCSF ChimeraX (Goddard et al., 2018).

### In vitro E2 discharge Assay for UBR5

We first loaded purified Ub onto the purified E2. Purified Cys-Ub was modified with Dylight800 maleimide at pH 7.4 by mixing them at a molar ratio of 1:1.5 and incubating the mixture for 1 h, then 5 mM DTT was added to quench the modification reaction. Excess Dylight800 was removed by a Superdex75 column. The fluorescently labeled Ub was incubated with 0.5 μM recombinant mouse E1 and 8 μM recombinant human E2D2 in 50 mM HEPES pH 7.5, 150 mM NaCl and 3 mM Mg-ATP for 30 min at room temperature to produce the Ub charged E2 (Ub–E2D2). The reaction was quenched with 20 mM EDTA. Next, the Ub–E2D2 at the final concentration of 0.5 μM was incubated with 0.75 μM purified wild-type or mutant UBR5 proteins for 3 min at room temperature. Each reaction was in 20 μl volume and the reaction buffer contained 50 mM HEPES pH 7.5 and 150 mM NaCl. The reactions were quenched with 4x SDS sample loading buffer and run on the 4-12% SDS-PAGE gels. The gels were scanned by fluorescent imaging in a ChemiDoc MP imager (Bio-Rad). The intensities of the Ub–E2D2 bands were quantified by ImageJ and normalized to the Ub–E2D2 only control.

### In vitro ubiquitylation assay

Our in vitro ubiquitylation assay solution contained 0.2 μM purified E1, 2 μM purified E2D2, 50 μM ubiquitin (R&D Systems), either 0.5 μM wildtype UBR5 or catalytically dead mutant UBR5(C2768S), and 45 μM His-tagged PEPCK1 in 40 μl reaction buffer (50 mM HEPES pH7.5, 150 mM NaCl, 10 mM MgCl2, 10 mM AMPPNP). The reaction system was incubated with 20 μl Ni-NTA agarose beads for 120 min at 37°C. Then, the beads were washed two times by incubating with the wash buffer (50 mM HEPES pH7.5, 150 mM NaCl, 50 mM imidazole). Finally, the ubiquitylated PEPCK1 was eluted with 500 mM imidazole in 50 mM HEPES pH7.5, 150 mM NaCl. The eluent was mixed with the reducing 2X SDS loading buffer, analyzed by SDS-PAGE, and subsequently immunoblotted with an FK2 antibody (Cayman Chemical, Cat # 14220).

### Generation of MDA-MB-231 cell lines expressing mutant UBR5 proteins

MDA-MB-231 cells were cultured in DMEM high glucose medium (Hyclone) supplemented with 10% (V/V) FBS and 1% penicillin and streptomycin at 37°C in a humidified 5% CO_2_ atmosphere. To construct the MDA-MB-231-shUBR5 stable cell line, lentiviruses were produced by cotransfection of 293T cells with psPAX2 (Addgene plasmid 12260) and pMD2.G (Addgene plasmid 12259) in a ratio of 4:3:1 (4/3/1μg for a 10 cm dish), and pLKO-shUBR5. Viral supernatants were collected at 24 h and 48 h post-transfection. MDA-MB-231 cells were infected with the shUBR5-expressing lentivirus and selected with puromycin. An empty vector was used as the negative control (shNC). To generate the UBR5 ubiquitin ligase mutant T1209M, R2293Q, Y2576A, F2732A, A2790W, V196K, L224K, R1854H, C2768S, NTD (876-2799 aa), and C2768A-reconstituted MDA-MB-231-shUBR5 cell line, cells were transfected with the respective plasmids using Lipofectamine 3000 reagent (Invitrogen, L3000008) according to the manufacturer’s protocol. Stable cell lines were selected using G418 (250 μg/mL) for 4 weeks and expression of the mutant proteins were confirmed by Western blotting.

### Western Blot Analysis

Cells were lysed in RIPA buffer (Thermo Scientific), and the lysates were centrifuged at 12,000 rpm for 30 min at 4°C. Supernatants were collected and protein concentration was quantified using a Bio-Rad protein assay (Bio-Rad, 5000006). Cell lysates were subjected to SDS-PAGE and transferred to PVDF membranes, followed by immunoblotting with antibodies against UBR5 (sc-515494, Santa Cruz Biotechnology), Parafibromin/CDC73 (A300-170A, Bethyl Laboratories), and GAPDH (sc-FL335, Santa Cruz Biotechnology). Detection and quantification of band intensities was conducted using Image J software. Bands were normalized to total protein by dividing the intensity of the band by the intensity of the total protein from the same sample on the same blot. The relative expression was calculated by dividing the normalized intensity of experimental groups by the normalized intensity of control group. The Western blot was done in duplicate.

## Acknowledgements

Cryo-EM datasets were collected at the David Van Andel Advanced Cryo-Electron Microscopy Suite in Van Andel Institute. We thank G. Zhao and X. Meng for facilitating data collection. We thank E. Finkin-Groner at Tri-Institutional Therapeutics Discovery Institute for the gift of human E2D2 expression plasmid and L. Cantley at Harvard Medical School for the gift of PEPCK1 expression plasmid. This work was supported by the US Department of Defense grant W81XWH2110261 (to X.M.), the US National Institutes of Health grant GM131754 (to H.L.), and the Van Andel Institute (to H.L.).

## Author contributions

F.W. Q.H, W.Z., X.M., G.L, and H.L. designed the research; F.W., Q.H., W.Z., Z.Y., and E. F.-G. performed research and analyzed the data; F.W. and H.L. wrote the first draft of the manuscript. All authors reviewed and revised the manuscript.

## Data availability

The cryo-EM 3D maps of the human UBR5 have been deposited at the Electron Microscopy Data Bank database with accession code EMD-27201 (dimer in C1 symmetry) EMD-27822 (dimer in C2 symmetry), and EMD-28646 (tetramer in C1 symmetry). The corresponding atomic models were deposited at the RCSB Protein Data Bank database with accession codes 8D4X,8E0Q, and 8EWI.

## REFERENCES

Adams, P.D., Afonine, P.V., Bunkóczi, G., Chen, V.B., Davis, I.W., Echols, N., Headd, J.J., Hung, L.-W., Kapral, G.J., and Grosse-Kunstleve, R.W. (2010). PHENIX: a comprehensive Python-based system for macromolecular structure solution. Acta Crystallographica Section D: Biological Crystallography 66, 213–221.

Ahel, J., Lehner, A., Vogel, A., Schleiffer, A., Meinhart, A., Haselbach, D., and Clausen, T. (2020). Moyamoya disease factor RNF213 is a giant E3 ligase with a dynein-like core and a distinct ubiquitin-transfer mechanism. Elife 9.

Berndsen, C.E., and Wolberger, C. (2014). New insights into ubiquitin E3 ligase mechanism. Nature structural & molecular biology 21, 301–307.

Callaghan, M.J., Russell, A.J., Woollatt, E., Sutherland, G.R., Sutherland, R.L., and Watts, C.K. (1998). Identification of a human HECT family protein with homology to the Drosophila tumor suppressor gene hyperplastic discs. Oncogene 17, 3479–3491.

Chen, C.K., Chan, N.L., and Wang, A.H. (2011). The many blades of the beta-propeller proteins: conserved but versatile. Trends Biochem Sci 36, 553–561.

Choi, W.S., Jeong, B.-C., Joo, Y.J., Lee, M.-R., Kim, J., Eck, M.J., and Song, H.K. (2010). Structural basis for the recognition of N-end rule substrates by the UBR box of ubiquitin ligases. Nature structural & molecular biology 17, 1175–1181.

Cojocaru, M., Bouchard, A., Cloutier, P., Cooper, J.J., Varzavand, K., Price, D.H., and Coulombe, B. (2011). Transcription factor IIS cooperates with the E3 ligase UBR5 to ubiquitinate the CDK9 subunit of the positive transcription elongation factor B. Journal of Biological Chemistry 286, 5012–5022.

Ding, F., Zhu, X., Song, X., Yuan, P., Ren, L., Chai, C., Zhou, W., and Li, X. (2020). UBR5 oncogene as an indicator of poor prognosis in gastric cancer. Experimental and therapeutic medicine 20, 1–1.

Duda, D.M., Scott, D.C., Calabrese, M.F., Zimmerman, E.S., Zheng, N., and Schulman, B.A. (2011). Structural regulation of cullin-RING ubiquitin ligase complexes. Current opinion in structural biology 21, 257–264.

Emsley, P., and Cowtan, K. (2004). Coot: model-building tools for molecular graphics. Acta crystallographica section D: biological crystallography 60, 2126–2132.

Gallagher, E., Gao, M., Liu, Y.-C., and Karin, M. (2006). Activation of the E3 ubiquitin ligase Itch through a phosphorylation-induced conformational change. Proceedings of the National Academy of Sciences 103, 1717–1722.

George, A.J., Hoffiz, Y.C., Charles, A.J., Zhu, Y., and Mabb, A.M. (2018). A Comprehensive Atlas of E3 Ubiquitin Ligase Mutations in Neurological Disorders. Front Genet 9, 29.

Goddard, T.D., Huang, C.C., Meng, E.C., Pettersen, E.F., Couch, G.S., Morris, J.H., and Ferrin, T.E. (2018). UCSF ChimeraX: Meeting modern challenges in visualization and analysis. Protein Science 27, 14–25.

Grabarczyk, D.B., Petrova, O.A., Deszcz, L., Kurzbauer, R., Murphy, P., Ahel, J., Vogel, A., Gogova, R., Faas, V., and Kordic, D. (2021). HUWE1 employs a giant substrate-binding ring to feed and regulate its HECT E3 domain. Nature Chemical Biology 17, 1084–1092.

Hay-Koren, A., Caspi, M., Zilberberg, A., and Rosin-Arbesfeld, R. (2011). The EDD E3 ubiquitin ligase ubiquitinates and up-regulates β-catenin. Molecular biology of the cell 22, 399–411.

Honda, Y., Tojo, M., Matsuzaki, K., Anan, T., Matsumoto, M., Ando, M., Saya, H., and Nakao, M. (2002). Cooperation of HECT-domain ubiquitin ligase hHYD and DNA topoisomerase II-binding protein for DNA damage response. Journal of Biological Chemistry 277, 3599–3605.

Horn-Ghetko, D., Krist, D.T., Prabu, J.R., Baek, K., Mulder, M.P., Klügel, M., Scott, D.C., Ovaa, H., Kleiger, G., and Schulman, B.A. (2021). Ubiquitin ligation to F-box protein targets by SCF–RBR E3– E3 super-assembly. Nature 590, 671–676.

Horn-Ghetko, D., and Schulman, B.A. (2022). New classes of E3 ligases illuminated by chemical probes. Curr Opin Struct Biol 73, 102341.

Hu, Q., Botuyan, M.V., Zhao, D., Cui, G., Mer, E., and Mer, G. (2021). Mechanisms of BRCA1-BARD1 nucleosome recognition and ubiquitylation. Nature 596, 438–443.

Hunkeler, M., Jin, C.Y., Ma, M.W., Monda, J.K., Overwijn, D., Bennett, E.J., and Fischer, E.S. (2021). Solenoid architecture of HUWE1 contributes to ligase activity and substrate recognition. Molecular cell 81, 3468–3480. e3467.

Jiang, H., He, X., Feng, D., Zhu, X., and Zheng, Y. (2015). RanGTP aids anaphase entry through Ubr5-mediated protein turnover. J Cell Biol 211, 7–18.

Jiang, W., Wang, S., Xiao, M., Lin, Y., Zhou, L., Lei, Q., Xiong, Y., Guan, K.L., and Zhao, S. (2011). Acetylation regulates gluconeogenesis by promoting PEPCK1 degradation via recruiting the UBR5 ubiquitin ligase. Mol Cell 43, 33–44.

Jumper, J., Evans, R., Pritzel, A., Green, T., Figurnov, M., Ronneberger, O., Tunyasuvunakool, K., Bates, R., Žídek, A., and Potapenko, A. (2021). Highly accurate protein structure prediction with AlphaFold. Nature 596, 583–589.

Kaisari, S., Miniowitz-Shemtov, S., Sitry-Shevah, D., Shomer, P., Kozlov, G., Gehring, K., and Hershko, A. (2022). Role of ubiquitin-protein ligase UBR5 in the disassembly of mitotic checkpoint complexes. Proc Natl Acad Sci U S A 119.

Kamadurai, H.B., Qiu, Y., Deng, A., Harrison, J.S., MacDonald, C., Actis, M., Rodrigues, P., Miller, D.J., Souphron, J., and Lewis, S.M. (2013). Mechanism of ubiquitin ligation and lysine prioritization by a HECT E3. Elife 2, e00828.

Kamadurai, H.B., Souphron, J., Scott, D.C., Duda, D.M., Miller, D.J., Stringer, D., Piper, R.C., and Schulman, B.A. (2009). Insights into ubiquitin transfer cascades from a structure of a UbcH5B~ ubiquitin-HECTNEDD4L complex. Molecular cell 36, 1095–1102.

Kim, J.G., Shin, H.C., Seo, T., Nawale, L., Han, G., Kim, B.Y., Kim, S.J., and Cha-Molstad, H. (2021). Signaling Pathways Regulated by UBR Box-Containing E3 Ligases. Int J Mol Sci 22.

Kostrhon, S., Prabu, J.R., Baek, K., Horn-Ghetko, D., von Gronau, S., Klügel, M., Basquin, J., Alpi, A.F., and Schulman, B.A. (2021). CUL5-ARIH2 E3-E3 ubiquitin ligase structure reveals cullin-specific NEDD8 activation. Nature Chemical Biology 17, 1075–1083.

Kozlov, G., Nguyen, L., Lin, T., De Crescenzo, G., Park, M., and Gehring, K. (2007). Structural basis of ubiquitin recognition by the ubiquitin-associated (UBA) domain of the ubiquitin ligase EDD. Journal of Biological Chemistry 282, 35787–35795.

Lechtenberg, B.C., Rajput, A., Sanishvili, R., Dobaczewska, M.K., Ware, C.F., Mace, P.D., and Riedl, S.J. (2016). Structure of a HOIP/E2~ ubiquitin complex reveals RBR E3 ligase mechanism and regulation. Nature 529, 546–550.

Li, J., Zhang, W., Gao, J., Du, M., Li, H., Li, M., Cong, H., Fang, Y., Liang, Y., and Zhao, D. (2021). E3 Ubiquitin Ligase UBR5 Promotes the Metastasis of Pancreatic Cancer via Destabilizing F-Actin Capping Protein CAPZA1. Frontiers in oncology 11, 677.

Li, W., Bengtson, M.H., Ulbrich, A., Matsuda, A., Reddy, V.A., Orth, A., Chanda, S.K., Batalov, S., and Joazeiro, C.A. (2008). Genome-wide and functional annotation of human E3 ubiquitin ligases identifies MULAN, a mitochondrial E3 that regulates the organelle’s dynamics and signaling. PLoS One 3, e1487.

Liao, L., Song, M., Li, X., Tang, L., Zhang, T., Zhang, L., Pan, Y., Chouchane, L., and Ma, X. (2017). E3 ubiquitin ligase UBR5 drives the growth and metastasis of triple-negative breast cancer. Cancer research 77, 2090–2101.

Lorenz, S. (2018). Structural mechanisms of HECT-type ubiquitin ligases. Biological chemistry 399, 127–145.

Lumpkin, R.J., Baker, R.W., Leschziner, A.E., and Komives, E.A. (2020). Structure and dynamics of the ASB9 CUL-RING E3 Ligase. Nat Commun 11, 2866.

Mansfield, E., Hersperger, E., Biggs, J., and Shearn, A. (1994). Genetic and molecular analysis of hyperplastic discs, a gene whose product is required for regulation of cell proliferation in Drosophila melanogaster imaginal discs and germ cells. Developmental biology 165, 507–526.

Maspero, E., Mari, S., Valentini, E., Musacchio, A., Fish, A., Pasqualato, S., and Polo, S. (2011). Structure of the HECT: ubiquitin complex and its role in ubiquitin chain elongation. EMBO reports 12, 342–349.

Maspero, E., Valentini, E., Mari, S., Cecatiello, V., Soffientini, P., Pasqualato, S., and Polo, S. (2013). Structure of a ubiquitin-loaded HECT ligase reveals the molecular basis for catalytic priming. Nature structural & molecular biology 20, 696–701.

Mastronarde, D.N. (2018). Advanced data acquisition from electron microscopes with SerialEM. Microscopy and Microanalysis 24, 864–865.

Matta-Camacho, E., Kozlov, G., Li, F.F., and Gehring, K. (2010). Structural basis of substrate recognition and specificity in the N-end rule pathway. Nat Struct Mol Biol 17, 1182–1187.

Matta-Camacho, E., Kozlov, G., Menade, M., and Gehring, K. (2012). Structure of the HECT C-lobe of the UBR5 E3 ubiquitin ligase. Acta Crystallogr Sect F Struct Biol Cryst Commun 68, 1158–1163.

Morreale, F.E., and Walden, H. (2016). Types of Ubiquitin Ligases. Cell 165, 248–248 e241.

Munoz-Escobar, J., Matta-Camacho, E., Cho, C., Kozlov, G., and Gehring, K. (2017). Bound Waters Mediate Binding of Diverse Substrates to a Ubiquitin Ligase. Structure 25, 719–729 e713.

Muñoz-Escobar, J., Matta-Camacho, E., Kozlov, G., and Gehring, K. (2015). The MLLE domain of the ubiquitin ligase UBR5 binds to its catalytic domain to regulate substrate binding. Journal of Biological Chemistry 290, 22841–22850.

Ogunjimi, A.A., Briant, D.J., Pece-Barbara, N., Le Roy, C., Di Guglielmo, G.M., Kavsak, P., Rasmussen, R.K., Seet, B.T., Sicheri, F., and Wrana, J.L. (2005). Regulation of Smurf2 ubiquitin ligase activity by anchoring the E2 to the HECT domain. Molecular cell 19, 297–308.

Ohtake, F., Tsuchiya, H., Saeki, Y., and Tanaka, K. (2018). K63 ubiquitylation triggers proteasomal degradation by seeding branched ubiquitin chains. Proceedings of the National Academy of Sciences 115, E1401–E1408.

Pan, M., Zheng, Q., Wang, T., Liang, L., Mao, J., Zuo, C., Ding, R., Ai, H., Xie, Y., Si, D., et al. (2021). Structural insights into Ubr1-mediated N-degron polyubiquitination. Nature 600, 334–338.

Petroski, M.D., and Deshaies, R.J. (2005). Function and regulation of cullin–RING ubiquitin ligases. Nature reviews Molecular cell biology 6, 9–20.

Pettersen, E.F., Goddard, T.D., Huang, C.C., Couch, G.S., Greenblatt, D.M., Meng, E.C., and Ferrin, T.E. (2004). UCSF Chimera—a visualization system for exploratory research and analysis. Journal of computational chemistry 25, 1605–1612.

Punjani, A., and Fleet, D.J. (2021). 3D variability analysis: Resolving continuous flexibility and discrete heterogeneity from single particle cryo-EM. Journal of Structural Biology 213, 107702.

Punjani, A., Rubinstein, J.L., Fleet, D.J., and Brubaker, M.A. (2017). cryoSPARC: algorithms for rapid unsupervised cryo-EM structure determination. Nature methods 14, 290–296.

Qiao, S., Langlois, C.R., Chrustowicz, J., Sherpa, D., Karayel, O., Hansen, F.M., Beier, V., von Gronau, S., Bollschweiler, D., Schafer, T., et al. (2020). Interconversion between Anticipatory and Active GID E3 Ubiquitin Ligase Conformations via Metabolically Driven Substrate Receptor Assembly. Mol Cell 77, 150–163 e159.

Rotin, D., and Kumar, S. (2009). Physiological functions of the HECT family of ubiquitin ligases. Nature reviews Molecular cell biology 10, 398–409.

Rutz, S., Kayagaki, N., Phung, Q.T., Eidenschenk, C., Noubade, R., Wang, X., Lesch, J., Lu, R., Newton, K., and Huang, O.W. (2015). Deubiquitinase DUBA is a post-translational brake on interleukin-17 production in T cells. Nature 518, 417–421.

Sanchez-Garcia, R., Gomez-Blanco, J., Cuervo, A., Carazo, J.M., Sorzano, C.O.S., and Vargas, J. (2021). DeepEMhancer: a deep learning solution for cryo-EM volume post-processing. Commun Biol 4, 874.

Sander, B., Xu, W., Eilers, M., Popov, N., and Lorenz, S. (2017). A conformational switch regulates the ubiquitin ligase HUWE1. Elife 6, e21036.

Scheres, S.H., and Chen, S. (2012). Prevention of overfitting in cryo-EM structure determination. Nature methods 9, 853–854.

Shah, S.S., and Kumar, S. (2021). Adaptors as the regulators of HECT ubiquitin ligases. Cell Death & Differentiation 28, 455–472.

Shakeel, S., Rajendra, E., Alcon, P., O’Reilly, F., Chorev, D.S., Maslen, S., Degliesposti, G., Russo, C.J., He, S., Hill, C.H., et al. (2019). Structure of the Fanconi anaemia monoubiquitin ligase complex. Nature 575, 234–237.

Shearer, R.F., Iconomou, M., Watts, C.K., and Saunders, D.N. (2015). Functional roles of the E3 ubiquitin ligase UBR5 in cancer. Molecular Cancer Research 13, 1523–1532.

Shen, Q., Qiu, Z., Wu, W., Zheng, J., and Jia, Z. (2018). Characterization of interaction and ubiquitylation of phosphoenolpyruvate carboxykinase by E3 ligase UBR5. Biol Open 7.

Sherpa, D., Chrustowicz, J., Qiao, S., Langlois, C.R., Hehl, L.A., Gottemukkala, K.V., Hansen, F.M., Karayel, O., von Gronau, S., Prabu, J.R., et al. (2021). GID E3 ligase supramolecular chelate assembly configures multipronged ubiquitin targeting of an oligomeric metabolic enzyme. Mol Cell 81, 2445–2459 e2413.

Sherpa, D., Chrustowicz, J., and Schulman, B.A. (2022). How the ends signal the end: Regulation by E3 ubiquitin ligases recognizing protein termini. Mol Cell 82, 1424–1438.

Singh, S., Ng, J., Nayak, D., and Sivaraman, J. (2019). Structural insights into a HECT-type E3 ligase AREL1 and its ubiquitylation activities in vitro. Journal of Biological Chemistry 294, 19934–19949.

Song, M., Yeku, O.O., Rafiq, S., Purdon, T., Dong, X., Zhu, L., Zhang, T., Wang, H., Yu, Z., and Mai, J. (2020). Tumor derived UBR5 promotes ovarian cancer growth and metastasis through inducing immunosuppressive macrophages. Nature communications 11, 1–16.

Sriram, S.M., and Kwon, Y.T. (2010). The molecular principles of N-end rule recognition. Nature structural & molecular biology 17, 1164–1165.

Walden, H., and Rittinger, K. (2018). RBR ligase–mediated ubiquitin transfer: a tale with many twists and turns. Nature structural & molecular biology 25, 440–445.

Wang, X., Singh, S., Jung, H.-Y., Yang, G., Jun, S., Sastry, K.J., and Park, J.-I. (2013). HIV-1 Vpr protein inhibits telomerase activity via the EDD-DDB1-VPRBP E3 ligase complex. Journal of Biological Chemistry 288, 15474–15480.

Weissman, A.M. (2001). Themes and variations on ubiquitylation. Nature reviews Molecular cell biology 2, 169–178.

Wiesner, S., Ogunjimi, A.A., Wang, H.-R., Rotin, D., Sicheri, F., Wrana, J.L., and Forman-Kay, J.D. (2007). Autoinhibition of the HECT-type ubiquitin ligase Smurf2 through its C2 domain. Cell 130, 651–662.

Williams, C.J., Headd, J.J., Moriarty, N.W., Prisant, M.G., Videau, L.L., Deis, L.N., Verma, V., Keedy, D.A., Hintze, B.J., Chen, V.B., et al. (2018). MolProbity: More and better reference data for improved all-atom structure validation. Protein Sci 27, 293–315.

Witus, S.R., Burrell, A.L., Farrell, D.P., Kang, J., Wang, M., Hansen, J.M., Pravat, A., Tuttle, L.M., Stewart, M.D., Brzovic, P.S., et al. (2021). BRCA1/BARD1 site-specific ubiquitylation of nucleosomal H2A is directed by BARD1. Nat Struct Mol Biol 28, 268–277.

Xiang, G., Wang, S., Chen, L., Song, M., Song, X., Wang, H., Zhou, P., Ma, X., and Yu, J. (2022). UBR5 targets tumor suppressor CDC73 proteolytically to promote aggressive breast cancer. Cell Death Dis 13, 451.

Yang, Y., Zhao, J., Mao, Y., Lin, G., Li, F., and Jiang, Z. (2020). UBR5 over-expression contributes to poor prognosis and tamoxifen resistance of ERa+breast cancer by stabilizing beta-catenin. Breast Cancer Res Treat 184, 699–710.

Yoshida, M., Yoshida, K., Kozlov, G., Lim, N.S., De Crescenzo, G., Pang, Z., Berlanga, J.J., Kahvejian, A., Gehring, K., and Wing, S.S. (2006). Poly (A) binding protein (PABP) homeostasis is mediated by the stability of its inhibitor, Paip2. The EMBO journal 25, 1934–1944.

Youkharibache, P., Veretnik, S., Li, Q., Stanek, K.A., Mura, C., and Bourne, P.E. (2019). The Small beta-Barrel Domain: A Survey-Based Structural Analysis. Structure 27, 6–26.

Zhang, T., Cronshaw, J., Kanu, N., Snijders, A.P., and Behrens, A. (2014). UBR5-mediated ubiquitylation of ATMIN is required for ionizing radiation-induced ATM signaling and function. Proceedings of the National Academy of Sciences 111, 12091–12096.

Zheng, S.Q., Palovcak, E., Armache, J.-P., Verba, K.A., Cheng, Y., and Agard, D.A. (2017). MotionCor2: anisotropic correction of beam-induced motion for improved cryo-electron microscopy. Nature methods 14, 331–332

